# Construction of a Heterologous Pathway in *Escherichia coli* for Terephthalate Assimilation

**DOI:** 10.1101/2025.02.25.640003

**Authors:** Tianyu Li, Nathan Crook

**Affiliations:** Department of Chemical Engineering, North Carolina State University, Raleigh, NC 27695, USA

**Keywords:** terephthalate, terephthalate acid, *Escherichia coli*, Polyethylene terephthalate, PET upcycling, bioremediation

## Abstract

Polyethylene terephthalate (PET) is extensively used in products such as packaging and textiles, and it remains a significant contributor to plastic and microplastic pollution. Effective valorization of PET waste requires not only depolymerizing it into its monomers but also utilizing the resulting products, ethylene glycol (EG) and terephthalic acid (TPA). While prior research has demonstrated that *Escherichia coli* can convert TPA into value-added compounds for PET upcycling, limited work has focused on its assimilation for cell growth. This study focuses on constructing a complete TPA metabolic pathway in *E. coli* comprising TPA uptake and its conversion to protocatechuate (PCA). We engineered *E. coli* JME3, which already possesses PCA catabolism, to utilize TPA as a sole carbon source by introducing a heterologous transporter TpaK and a PCA-synthesizing enzyme cascade. The strain next underwent adaptive laboratory evolution (ALE) and expression tuning to improve growth performance. The final strain achieved a maximum growth rate of 0.25 h⁻¹, demonstrating efficient TPA assimilation. These results expand the range of value-added compounds derived from PET upcycling and establish *E. coli* as a platform for coupling PET depolymerization with microbial growth, enabling high-throughput optimization of enzyme activity and PET biodegradation conditions.

## Introduction

Polyethylene terephthalate (PET), a widely used plastic, had a global market volume of 25.47 million metric tons in 2022 and is projected to reach 35.70 million metric tons by 2030 (“PET market volume worldwide 2015-2030,” 2024). However, its persistence in the environment as plastic and contribution to microplastic pollution has raised significant environmental concerns. In response, emerging research has focused on expressing PET-hydrolyzing enzymes (PHEs) in various microorganisms to degrade PET into its monomers, offering a bioremediation approach to address PET waste pollution. These strategies include intracellular enzyme expression followed by cell lysis or secretion (Shi et al., 2021), as well as surface display mechanisms (Chen et al., 2020; Gercke et al., 2021, 2021; Li et al., 2023; Zhu et al., 2022) to enhance PET breakdown into its monomers: ethylene glycol (EG) and terephthalate (TPA). However, achieving closed-loop recycling requires not only PET depolymerization but also the sustainable repurposing of its degradation products.

The microbial assimilation and valorization of EG have been extensively studied in species such as *Escherichia coli* (Kim et al., 2019; Pandit et al., 2021) and *Pseudomonas putida* (Franden et al., 2018; Li et al., 2019; Mückschel et al., 2012; Werner et al., 2021), leading to the production of value-added compounds like glycolic acid. In addition, TPA holds great recycling potential due to its molecular complexity and ability to serve as a precursor for high-value aromatic products. On a per-mole basis, TPA contains six more carbon molecules than EG, reinforcing its potential as a valuable substrate for bioconversion. Although natural TPA catabolism has been well characterized in environmental microorganisms such as *Ideonella sakaiensis*, *Comamonas* spp. E6, and *Rhodococcus jostii* RHA1 (Hara et al., 2007; Sasoh et al., 2006; Yoshida et al., 2016), its heterologous assimilation in model organisms remains underexplored. Among model organisms, *Pseudomonas putida* has been the most extensively engineered species for TPA assimilation, with significant progress demonstrated by Werner et al (Werner et al., 2021). In contrast, *E. coli* faces challenges in TPA catabolism due to the absence of a native TPA transporter and its limited metabolic capacity for aromatic compounds, particularly its inability to metabolize protocatechuic acid (PCA), a key intermediate in TPA degradation. Nevertheless, given *E. coli*’s widespread use in synthetic biology and industrial applications, enabling TPA assimilation in this host would expand its utility in plastic bioremediation and aromatic compound metabolism.

Efforts to upcycle TPA using *E. coli* have primarily focused on converting TPA into value-added products such as gallic acid, pyrogallol, and vanillic acid through the introduction of heterologous enzymes (Kim et al., 2019; Li et al., 2024; Sadler and Wallace, 2021) rather than enabling cell growth. To circumvent the challenge of active TPA transport, researchers have employed strategies such as enhancing cell membrane permeability with chemical additives like *n*-butanol (Sadler and Wallace, 2021) or implementing genetic modifications on membrane proteins (Li et al., 2024). Additionally, approaches involving high-density *E. coli* cultures coupled with cell autolysis have been used to release extracellular enzymes, facilitating TPA conversion (Kim et al., 2019; Valenzuela-Ortega et al., 2023).

Our study directly addresses the integration of TPA assimilation pathways within *E. coli*. Adopting an approach similar to Werner et al. (Werner et al., 2021), we incorporate transporters and enzyme cascades—including dioxygenases, reductases, and dehydrogenases—from *Rhodococcus jostii* and *Comamonas* spp. E6 into *E. coli* plasmids. We employ an engineered *E. coli* strain, JME3, which is capable of metabolizing protocatechuic acid (PCA) (Clarkson et al., 2017) as the chassis for expressing heterologous enzymes and transporters for TPA assimilation. Notably, we demonstrate the functionality of the TpaK transporter from *Rhodococcus jostii* in *E. coli*, enabling efficient TPA uptake without reliance on external treatments that compromise cellular integrity. This capability is validated by cell growth in minimal medium with TPA as the sole carbon source.

To further enhance strain performance, we apply adaptive laboratory evolution (ALE), followed by whole-genome sequencing to identify genetic adaptations, including changes in plasmid copy number, that contribute to improved growth. Additionally, We further optimize gene expression by fine-tuning transcription through an ribosome-binding site (RBS) library-based approach. These advancements expand the potential for TPA or PET valorization beyond aromatic compounds to a broader range of biosynthesized products in *E. coli*.

## Materials and methods

### Strains and culture conditions

Chemically competent *Escherichia coli* NEB5α (NEB #C2987) was used for all plasmid cloning. *Pseudomonas putida* KT2440 was purchased from ATCC. The *E. coli* derivative AG978 (BW25113 Δ*ompT::pcaHGBDC ΔpflB::pcaIJFK*), its evolved variant JME3 (Clarkson et al., 2017), and *Rhodococcus jostii* RHA1 were generously provided by Dr. Adam Guss.

All liquid cultures were incubated at 37°C with shaking at 250 rpm. For routine cultivation, *E. coli* and its derivatives were grown in Miller’s LB broth (BD B244610). To assess cell growth, strains were cultivated in freshly prepared minimal media. Minimal media referred to here as M9 minimal medium, was formulated with 3 g/L KH₂PO₄, 6.78 g/L Na₂HPO₄, 0.5 g/L NaCl, 1 g/L NH₄Cl, 2 mM MgSO₄, 0.1 mM CaCl₂, and 18 μM FeSO₄, supplemented with various carbon sources, including 0.4% glucose, 10 mM TPA, or 1 g/L PCA. When required, 1 mM isopropyl ß-D-1-thiogalactopyranoside (IPTG) was added for induction. Antibiotics were included as needed to maintain plasmids: 35 μg/mL chloramphenicol (Cm) or 50 μg/mL kanamycin (Kan).

TPA stock solutions (100 mM) were prepared by dissolving terephthalic acid (TPA, CAS: 100-21-0) in deionized (DI) water, adding 350 mM NaOH to ensure complete dissolution and establish a high pH environment (pH = 13). The solution was then filtered through a 0.2 μm membrane for sterilization, yielding TPA minimal media with a final pH of approximately 7.5. Additionally, a similar stock solution (100mM) was prepared using disodium terephthalate (Na_2_TP, CAS: 10028-70-3) dissolved in DI water, with a trace amount of NaOH added to adjust the pH to 7. To avoid confusion, the minimal medium prepared in this way will be referred to as Na_2_TP minimal media.

### Plasmid and strain construction

Plasmids were constructed using Gibson Assembly with the NEBuilder HiFi DNA Assembly Master Mix (NEB #E2621). Operons encoding the enzyme cascade (*tphA2A3BA1*) from *Comamonas* sp. E6—including dioxygenase, reductase, and dehydrogenase—were synthesized by Twist Bioscience and cloned into a pET28 backbone (a gift from Dr. Chase Beisel) under the *Ptrc* promoter with a kanamycin resistance marker. Transporter genes, including *tpaK* from *Rhodococcus jostii* RHA1 (amplified using Q5 Hot Start High-Fidelity 2X Master Mix, NEB #M0494) and *tphC:tpiBA* (synthesized as an operon by Twist Bioscience, shown in Supplementary File 1.) from *Comamonas* sp. E6, were cloned into a pACYC184 backbone (TelesisBio) under the *Ptrc* promoter with a chloramphenicol resistance marker. Ribosome binding sites (RBSs) were optimized for varying translation initiation rates (TIRs) using De Novo DNA to balance transporter and enzyme expression.

Constructed plasmids were verified via colony PCR using Phire Plant Direct PCR Master Mix (Thermo Scientific, Cat. F160S) with primer pairs synthesized by Eurofins Genomics. Coding regions were confirmed by Sanger sequencing (GENEWIZ, Inc.), followed by whole-plasmid sequencing (Plasmidsaurus, Inc.). Detailed plasmid maps are provided in Supplementary File 3, and all primer sequences are listed in Supplementary File 1.

Plasmids were extracted using the Zyppy Plasmid Miniprep Kit (Zymo Research, D4019) and retransformed into *E. coli* AG978 and JME3. Chemically competent cells were prepared and transformed following *E. coli* protocols adapted from Chung and Miller (Chung and Miller, 1993). Each construct was preserved as frozen stocks by mixing 500 μL of overnight culture with 500 μL of 50% (v/v) glycerol in deionized (DI) water and stored at −80°C.

#### Growth Assays

Strains were revived from frozen stocks onto LB agar plates with the appropriate antibiotics and incubated in the 37°C standing incubator overnight. Single colonies were inoculated into LB broth and grown overnight in the shaking incubator at 37°C, 250 rpm. Cultures were then sub-inoculated into 0.4% glucose minimal medium supplemented with 1 mM IPTG, and relevant antibiotics. These cultures were grown to the stationary phase to acclimate to minimal medium and allow enzyme expression before being diluted 1:100 into fresh minimal medium with 0.4% glucose for subculturing.

At the early exponential phase (OD600 = 0.2–0.4), 1 mM IPTG was added again to induce transporter and enzyme expression. *E. coli* derivatives were harvested at the stationary phase (∼5 hours post-induction), washed three times with PBS, and resuspended in fresh TPA or Na_2_TP minimal medium. Final cultures were adjusted to an initial OD_600_ of 0.05 and incubated with shaking.

#### Adaptive laboratory evolution experiments

Adaptive laboratory evolution (ALE) experiments were initiated by subculturing cultures with observed growth from the growth assay into freshly prepared TPA minimal medium at a 1:100 dilution. Cultures were passaged at recorded intervals after measuring the final optical density, and frozen stocks were saved after each passage. This process was repeated for nine passages, yielding ten distinct populations for each strain (Supplementary File 2). The 1st, 5th, and 10th populations were subjected to serial dilution and plated onto LB agar plates to isolate individual colonies from the frozen stocks. Ten individual colonies from these three populations were then cultured in TPA minimal medium and stored as frozen stocks. These isolates and populations were subsequently analyzed through growth curve measurements in both TPA and Na_2_TP minimal media using a microplate reader and genome sequencing for further characterization.

#### Whole-genome sequencing and analysis

Genomic DNA was extracted from selected isolates grown to stationary phase in minimal medium with TPA using the Quick-DNA Fungal/Bacterial Miniprep Kit (Zymo Research, D6005). Genome samples were sequenced using Illumina NovaSeq 6000 sequencer by SeqCenter. The wild-type genome of *E. coli* JME3 was sequenced to establish mutation baselines. All sequences have been deposited in the Sequence Read Archive under accession number PRJNA1222917. Sequencing data were analyzed using *breseq* (Barrick et al., 2014) with the following references: *Escherichia coli* BW25113 (CP009273) and two plasmids (plaTL055, plaTL057).

#### RBS Library Construction

The pET28 plasmid was used as a backbone to express transporter and enzyme cascades as operons under the *Ptrc* promoter. The ribosome binding site (RBS) library was designed using De Novo DNA for *E. coli* JME3 and is shown in Supplementary File 1. The vector backbones were amplified using Q5 Hot Start High-Fidelity 2X Master Mix (NEB #M0494), while the RBS library was generated through PCR amplification with NEBNext® Ultra™ II Q5® Master Mix (NEB #M0544) to improve the uniformity of library amplification. Inserts and vector backbones were assembled using Gibson Assembly (NEBuilder HiFi DNA Assembly Master Mix, NEB #E2621) into plaTL097, purified using the DNA Clean & Concentrator Kit (Zymo, #D4004), and transformed into NEB10β electrocompetent *E. coli* (NEB #C3020) for optimal transformation efficiency.

To evaluate the library, 20 μL of recovered culture was serially diluted and spot-plated on LB agar plates with kanamycin to assess library size. Ten colonies from the overnight culture plates were inoculated, and plasmids were extracted to confirm the RBS library by sequencing (Plasmidsaurus). The remaining cells were cultured in 100 mL LB broth with kanamycin and incubated at 26°C, 250 rpm, until an OD600 of approximately 0.6 was reached. Low-temperature amplification was used to minimize amplification bias. Plasmids were then extracted using the Zyppy Plasmid Midiprep Kit (Zymo Research, #D4200) and transformed into *E. coli* JME3 by electroporation.

Recovered cells were cultured overnight in LB with kanamycin at 26°C, 250 rpm. Afterward, cells were washed with PBS and resuspended in 100 mL minimal medium with 10 mM Na₂TP for *E. coli* JME3, with an initial OD600 of 0.05 for screening. Cultures from library screening were harvested at the stationary phase, and plasmids were extracted for sequencing by Plasmidsaurus (Supplementary File 3). The raw data were analyzed using the DADA2 package in R (Callahan et al., 2016). The DADA2 analysis was performed using custom code, which was adapted for primer removal, filtering, error learning, denoising, and clustering of amplicon sequences (Supplementary File 3). The remaining library cultures were plated on 10 mM Na₂TP minimal agar plates to isolate individual variants. Ten colonies from each library were picked, subcultured in Na₂TP minimal medium, and stored as frozen stocks. The growth of these isolates was further assessed using a microplate reader. Plasmids from selected isolates were then extracted using the Zyppy Plasmid Miniprep Kit for additional sequencing and analysis.

#### Evaluation of Growth in Microtiter Plates

Frozen stocks of evolved populations were thawed, and a small aliquot was diluted 1:100 into fresh TPA minimal medium. The diluted culture was then divided into six technical replicates of 200 µL each for direct population growth measurement using a microplate reader. Unlike populations, individual variants were pre-cultured in 96-well microplates (Genesee) with 200 µL of TPA minimal medium and grown to the stationary phase. Endpoint OD600 measurements were taken using a microplate reader, and cultures were normalized to an OD600 of 0.01 in fresh 200 µL TPA minimal medium in 96-well microplates.

Growth curves were recorded using a Tecan Sunrise microplate reader, with absorbance at 600 nm measured every 10 minutes under continuous orbital shaking over approximately 72 hours. Intrinsic growth rates (µ), carrying capacities (K), and the time at which the population density reaches half of its carrying capacity (t_mid_) were determined by fitting the data to a logistic growth equation using the *growthcurver* package in R (Sprouffske and Wagner, 2016). Whole-plate growth plots were also generated using *growthcurver* for visualization and analysis. All growth data is documented in Supplementary File 4.

## Results

### Development of TPA Catabolism in *E. coli*

To establish a functional TPA assimilation pathway, we began with *E. coli* strains AG978 and JME3 (Clarkson et al., 2017). *E. coli* AG978 is a derivative of *E. coli* BW25113 which carries the PCA 3,4-cleavage pathway (*pcaHGBCDIJF*) and the PCA transporter gene *pcaK* from *Pseudomonas putida* KT2440. These genes were integrated into the *ompT* and *pflB* loci of *E. coli* BW25113 (ΔompT::pcaHGBDC ΔpflB::pcaIJFK). JME3, an evolved variant of AG978, exhibits enhanced carrying capacity and robust growth rates on PCA-containing media, as reported by Clarkson et al. (Clarkson et al., 2017).

We first compared the growth of *E. coli* AG978 and JME3 in PCA minimal medium (1 g/L) to that of *E. coli* BW25113 at 37°C. Growth curves in PCA minimal medium are shown in Figure 1, with growth in 0.4% glucose minimal medium provided in Supplementary Figure S1. As it has not been engineered for growth on PCA, *E. coli* BW25113 exhibited no growth in PCA medium. *E. coli* AG978 showed no detectable growth within 25 hours, but exhibited growth after 2–3 days upon extended incubation (data not shown). In contrast, *E. coli* JME3 exhibited a intrinsic growth rate of 0.60 ±0.03 h⁻¹, indicating its ability to effectively assimilate PCA. JME3 was therefore selected as the chassis strain for further pathway development.

**Figure 1.**
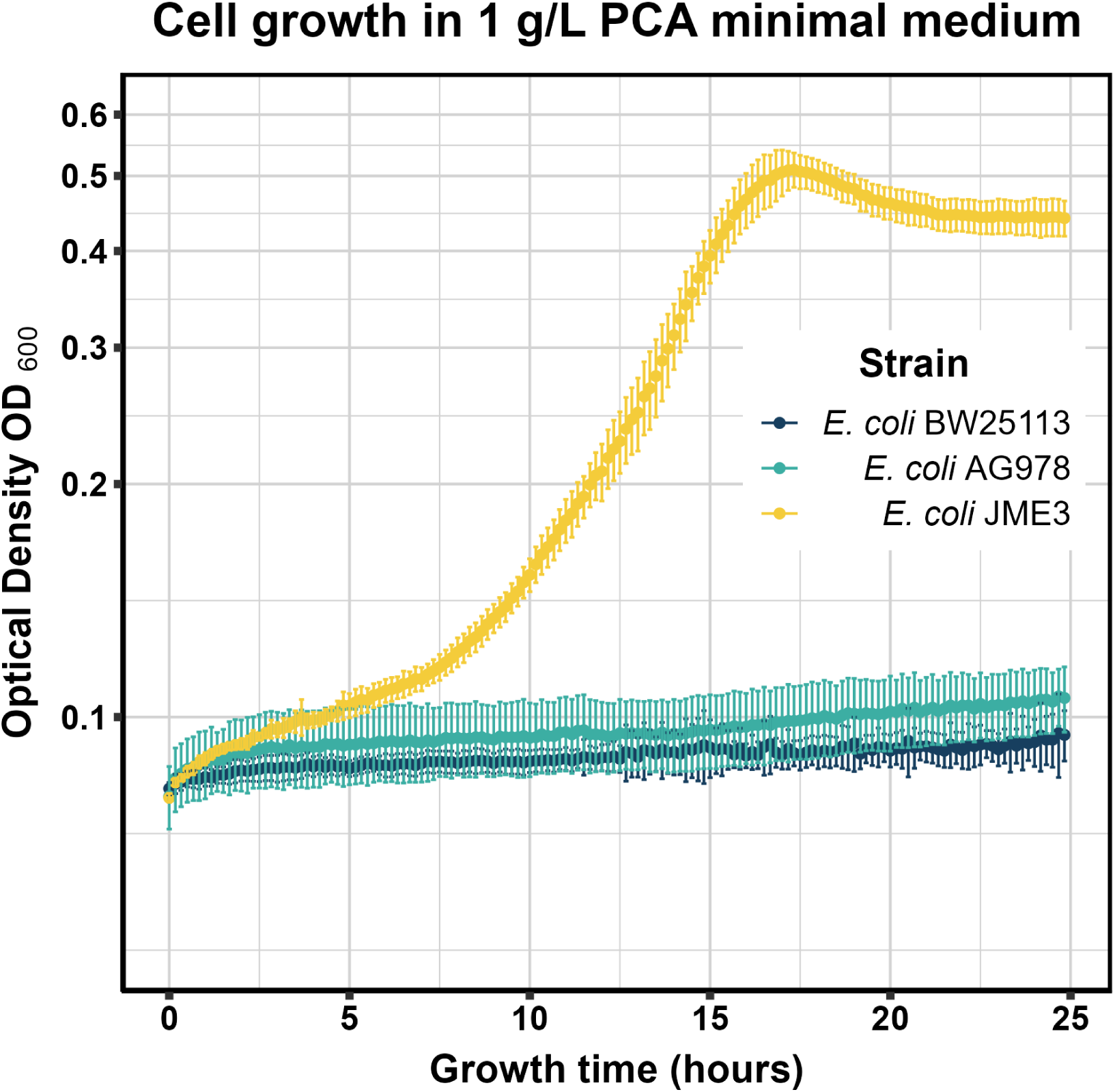
Growth curves of *E. coli* BW25113, AG978, and JME3 in minimal medium containing 1 g/L protocatechuic acid as the sole carbon source. Results are presented as mean values, with error bars representing the standard deviation from six technical replicates for each strain.The y-axis is plotted on a logarithmic scale.

**Figure 2.**
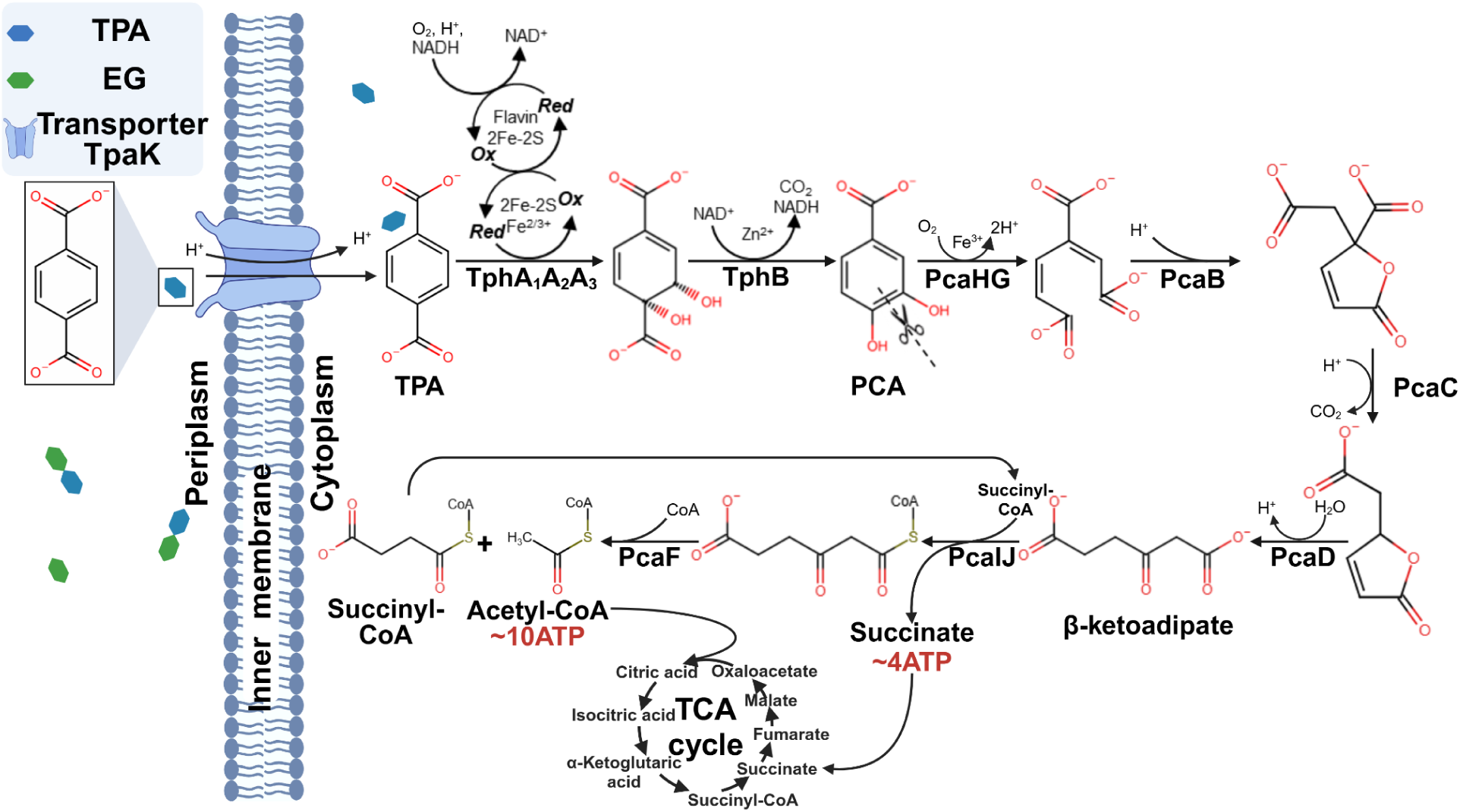
Schematic of TPA metabolism pathway. This schematic depicts TPA uptake via the TpaK transporter and its sequential conversion to PCA and β-ketoadipate through key enzymatic steps. Metabolic intermediates and their integration into the tricarboxylic acid (TCA) cycle are highlighted. The ATP-equivalent energy yield from acetyl-CoA and succinate oxidation is also indicated.

Following the approach of Werner et al. (2021), TPA transporters and enzyme cascades from *Comamonas* sp. E6 and *R. jostii* RHA1 were considered for this study. The enzyme cascade *tphA2A3BA1* from *Comamonas* sp. E6, which converts TPA to PCA, was selected due to its successful application in *E. coli* for TPA valorization (Kim et al., 2019; Li et al., 2024; Sadler and Wallace, 2021). To facilitate TPA uptake, we tested the ATP-binding cassette transporter TpaK from *R. jostii* RHA1 and the tripartite transporter TphC:TpiBA from *Comamonas* sp. E6.

TPA transporters (*tpaK* and *tphC*:*tpiBA*) were expressed in constructs plaTL055 and plaTL056, respectively, while the enzyme cascade (*tphA2A3BA1*) was expressed in construct plaTL057. The enzyme cascade and one transporter were transformed into JME3, generating strains J055&057 (with the *tpaK* transporter) and J056&057 (with *tphC:tpiBA*) for further testing. Additionally, empty plasmids plaTL062 and plaTL063, which contained the same vector backbones as plaTL055/56 and plaTL057, respectively, but lacking coding sequences, were constructed and transformed into JME3, yielding the control strain J062&063.

Strains J055&057, J056&057, and J062&063 were initially subcultured in M9 minimal medium containing 0.4% glucose and subcultured into TPA minimal medium to acclimate to the minimal media environment. Given that PET-hydrolyzing enzymes such as leaf-branch compost cutinase (LCC) (Chen et al., 2023) and PETase from *Ideonella sakaiensis* (Lee et al., 2024) function optimally under slightly alkaline conditions (pH 8.0–9.0), TPA minimal media was prepared with an elevated pH. TPA stock solutions (100 mM) were prepared using 350 mM NaOH and then diluted 10-fold to prepare the medium, resulting in a final medium containing an excess of 15 mM NaOH after neutralizing TPA-derived protons. While this pH adjustment slightly suppressed growth, the effect was not significant (Supplemental Figure S2).

After 5 days of cultivation, growth was observed in J055&057 but not in J056&057 or the control strain J062&063, confirming the functionality of the TpaK transporter and TphA2A3BA1 enzyme cascade in *E. coli*. However, the long lag phase and low growth rate of this engineered strain, compared with JME3, prompted us to perform adaptive laboratory evolution to enhance its ability to grow on TPA.

### Adaptive Laboratory Evolution Enhances Growth on TPA

To initiate adaptive laboratory evolution, turbid cultures of J055&057 were subcultured into the fresh TPA minimal medium, with the goal of generating isolates with improved growth rates and reduced lag phase. Cultures were passaged upon reaching stationary phase a total of nine times (∼64 generations), yielding ten evolved populations, including the first population that exhibited growth. Details of the passaging process, along with the final culture optical density and timeline, are shown in Supplementary Figure S3. The 1st, 5th, and 10th populations (P1, P5, and P10) were selected for further analysis and their growth on TPA minimal medium was monitored using a microplate reader. Growth curves of these populations are shown in Figure 3. All three populations showed similar intrinsic growth rate. However, J055&057 P10 exhibited a significantly shorter lag phase than P1 and P5, suggesting that adaptive mutations enhanced its fitness in TPA minimal medium.

**Figure 3.**
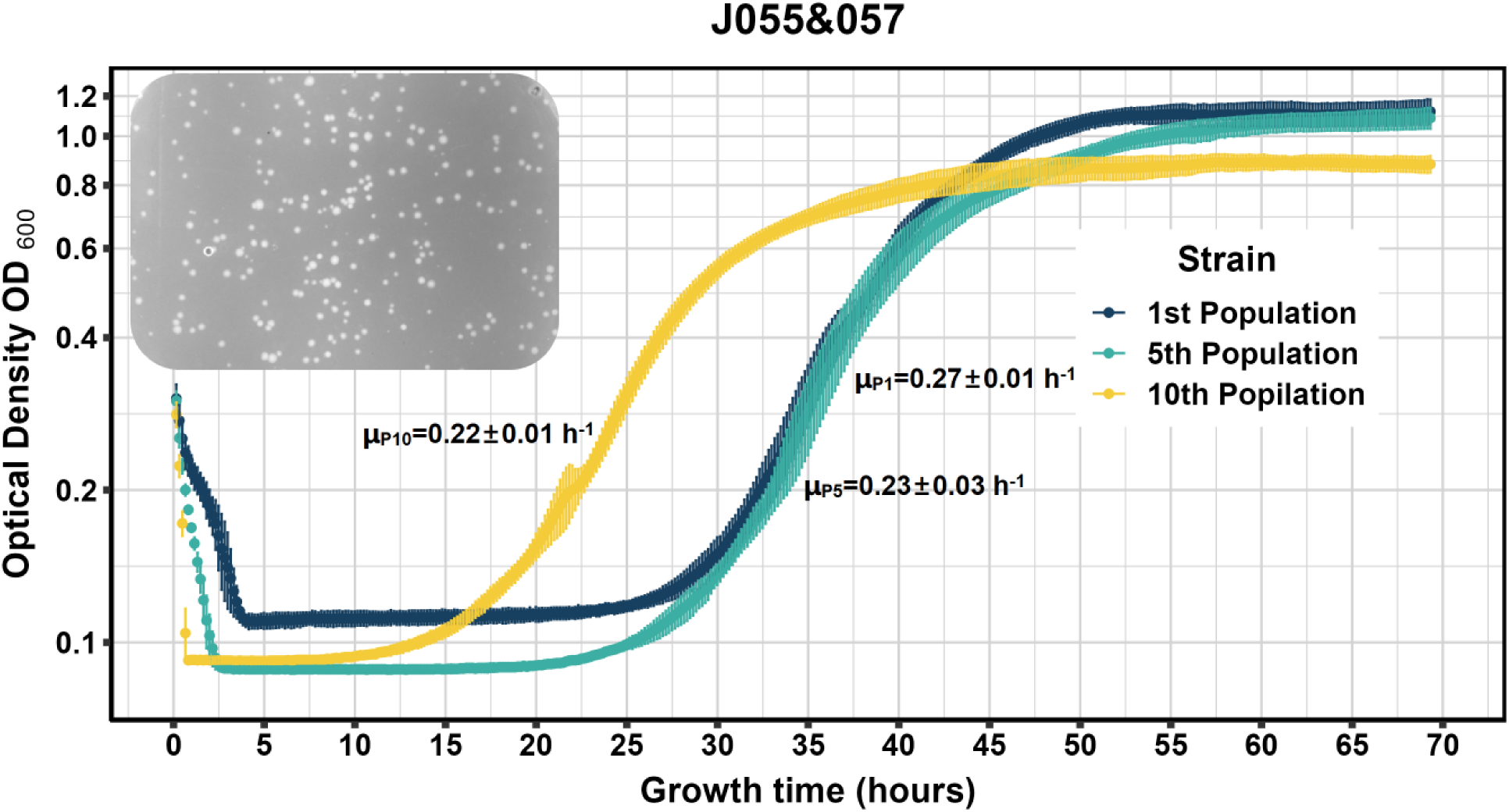
Growth curves of J055&057 P1, P5, and P10 in TPA minimal medium and their colony morphology. Results are presented as mean values, with error bars representing the standard deviation from six technical replicates. The y-axis is plotted on a logarithmic scale. Additionally, the calculated intrinsic growth rates with standard deviation from six technical replicates are included. A representative image of colonies on agar plates is shown in the top left corner, highlighting differences in colony size.

P1, P5, and P10 were spread onto agar plates to isolate individual variants. Notable differences in colony morphology were observed, particularly in P5 and P10, where colonies appeared to exhibit either a “large” or a “small” size (Figure 3). Ten variants were isolated from each population. Among them, isolates 1–5 from P5 and P10 exhibited the “small” colony morphology, whereas isolates 6–10 exhibited the “large” morphology. All variants were then subjected to individual growth curve measurements in two different types of media, either containing NaOH-neutralized TPA (as above) or in media containing an equivalent concentration of Na₂TP (Figure 4).

**Figure 4.**
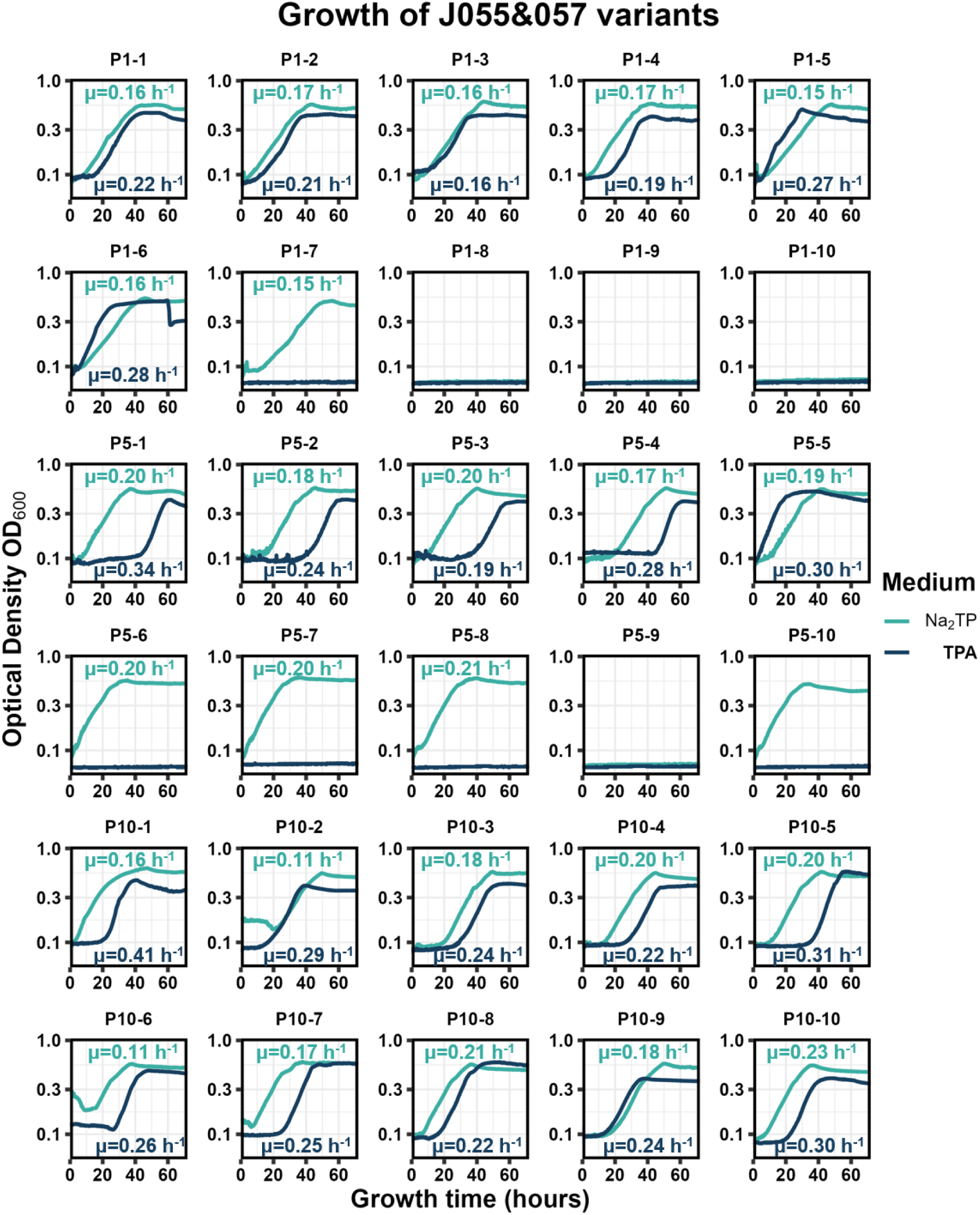
Growth curves of J055&057 variants isolated from P1, P5, and P10 in TPA and Na₂TP minimal media. The numbers following the dash represent individual isolate numbers. The y-axis is plotted on a logarithmic scale. Intrinsic growth rates are noted on the figure, with colors corresponding to different media conditions for clarity.

Most isolates exhibited distinct growth patterns in the two media types, most notably isolates P1-7 and P5-6 to P5-10, which grew in Na₂TP minimal medium but were inhibited in TPA minimal medium. These observations suggest that several variants were sensitive to high pH. In general, most variants displayed either a shorter lag phase or a higher carrying capacity in Na₂TP minimal medium compared to TPA minimal medium, except for P1-6 and P5-5, which performed better in TPA minimal medium. Overall, this suggests that J055&057 evolved not only for enhanced growth in TPA minimal medium but also for improved tolerance to elevated pH, which may otherwise repress cell growth. Taken together, these results provide evidence of adaptation and evolutionary changes over approximately 64 generations, resulting in distinct growth phenotypes. This motivated further investigation of several variants to analyze their genomes and identify the mutations contributing to their enhanced fitness.

### Whole-genome sequencing results

Three variants from each population— P1 (P1-2, P1-3, P1-6), P5 (P5-2, P5-3, P5-7), and P10 (P10-1, P10-5, P10-7)— along with the original JME3 strain as a control, were selected for whole-genome sequencing. As the complete genome sequence of JME3 was unavailable, whole-genome sequencing data for this strain was assembled using SPAdes (Bankevich et al., 2012) and annotated with Prokka (Seemann, 2014). Two contigs containing Δ*ompT::pcaHGBDC* and Δ*pflB::pcaIJFK* were identified via this procedure (Supplementary File 3). Due to the incomplete nature of the *de novo* assembly, the complete genome of BW25113, along with additional known contigs, and two plasmids plaTL055, and plaTL057, was used to identify *de novo* mutations in the evolved variants using *breseq* (Barrick et al., 2014).

The mutations we identified are shown in Figure 5 and Supplementary File 2. We observed some interesting adaptive patterns early in the evolutionary process. By analyzing genomic regions missing read coverage, we identified a 34–36 kb deletion between the *wsz* and *insF1* genes in three sequenced isolates (P1-2, P1-3, and P1-6) from P1. This deletion was absent in the sequenced variants from P5 and P10. The deleted region includes genes involved in capsule formation, envelope structure, nucleotide metabolism, DNA repair and alkylation response, regulatory functions, and transporters/efflux pumps. The loss of this deletion in further-evolved populations suggests a general disadvantage in strains containing it. We additionally observed a K17E mutation to *garL*, which encodes a 5-keto-4-deoxy-D-glucarate aldolase that splits 5-keto-4-deoxy-D-glucarate aldolase into pyruvate and tartronate semialdehyde (TSA) (Levy et al., 2025). This mutation is predicted to lie distal to the active site on a surface exposed residue, and therefore its contribution to protein function is unknown. The G11A mutation to *ybdH* (encoding hydroxycarboxylate dehydrogenase A) in P1-2 may alter its dehydrogenase function. YbdH catalyzes the NADPH-dependent reduction of 2-oxobutanoate and 2-oxoglutarate to 2-hydroxybutanoate and 2-hydroxyglutarate, respectively (Sévin et al., 2017). Since 2-oxoglutarate (α-ketoglutarate) is a key TCA cycle intermediate, its conversion to the toxic 2-hydroxyglutarate by YbdH could impair energy production and induce cellular stress (Sévin et al., 2017). Meanwhile, 2-oxobutanoate, another toxic intermediate, might be detoxified by pyruvate formate-lyase (PflB) along with putative pyruvate formate-lyase (TdcE), generating propionyl-CoA or propionate (Fang et al., 2021). However, in JME3, *pflB* was replaced with *pcaIJFK* for PCA assimilation, potentially disrupting 2-oxobutanoate detoxification. The *ybdH* (G11A) mutation may compensate by either reducing YbdH activity to preserve 2-oxoglutarate flux in the TCA cycle or enhancing its role in detoxifying 2-oxobutanoate, thereby mitigating toxicity in the absence of *pflB*.

**Figure 5.**
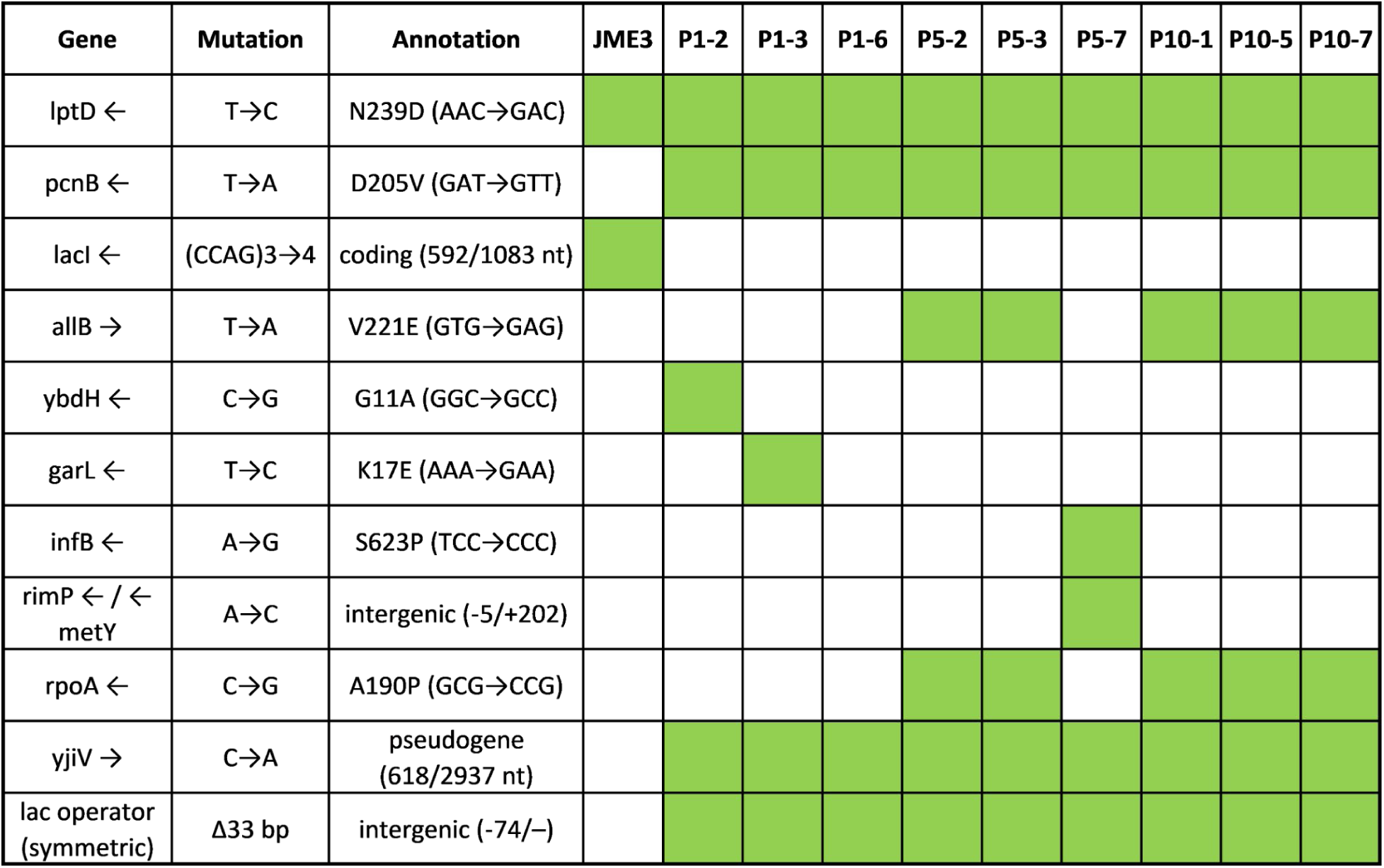
Summary of de novo mutations. Mutations detected are highlighted in green. Mutations in JME3 previously identified by Clarkson et al. are not shown. Additional details are provided in Supplementary File 2.

We also identified several mutations that potentially have more global effects on cellular physiology. First, we observed an N239D mutation in *lptD* in JME3, which facilitates lipopolysaccharide transport to the bacterial surface (Li et al., 2015). This mutation, not reported by Clarkson, *et al*, occurs in LptD’s extracellular loop 1, which could affect JME3’s envelope integrity, potentially altering stress responses and transcriptional regulation under PCA metabolism. We also observed an A190P mutation in *rpoA* (encoding the alpha subunit of RNA polymerase) in most isolates from P5 and P10. The isolate without an *rpoA* mutation (P5-7) instead contained an S623P mutation in *infB* (encoding translation initiation factor IF-2) and a mutation to the region upstream of *rimP*, encoding ribosome assembly factor P. This mutation is predicted to confer a reduced *rimP* translation rate (8.62 fold), which could result in a lower number of functional ribosomes available for translation. Coincident in strains with the *rpoA* mutation, we observed a V221E mutation in *allB*, which encodes an allantoinase for utilizing allantoin as nitrogen source under anaerobic conditions (Rodionova et al., 2023). As this mutation occurs away from AllB’s active site on a solvent exposed location, the rationale for this mutation is not known and it may be a hitchhiker in these strains. We expect that these mutations to *rpoA*, *infB*, and *rimP* could induce global changes to gene expression or cell growth.

Most prominently, we observed mutations consistent with an upregulation of PCA and TPA catabolic genes. Related to PCA catabolism, we identified that a tandem repeat at position 592 in the *lacI* gene (CCAG) increased from 3 to 4 copies in JME3, thus introducing a premature stop codon. This mutation, absent from previous sequencing results by Clarkson et al., corresponds to the upregulation of *pcaHGBDC* observed in other variants studied by them (Clarkson et al., 2017). However, this *lacI* mutation was not detected in the evolved isolates, likely due to higher read coverage of the intact *lacI* sequence from plaTL057 masking the chromosomal mutation. Instead, all isolates exhibited a 33-bp deletion upstream of the *pcaHGBDC* operon, which corresponds to the part of the symmetric lac operator known for tightly binding the *lac* repressor (Sadler et al., 1983). However, analysis of new read junctions revealed a 73.6% frequency of the same 33-bp deletion, suggesting that the JME3 strain used for sequencing had already partially evolved to eliminate this operator. Given that J055&057 carry an intact *lacI* gene in plaTL057, thus compensating for the chromosomal *lacI* mutation, this symmetric lac operator deletion likely alleviated repression, enhanced *pcaHGBDC* expression, and promoted better growth on TPA for the J055&057 variants.

Interestingly, no significant single nucleotide polymorphisms were observed in *plaTL055* or *plaTL057*. This may be due to the presence of multiple plasmid copies per cell, which would dilute the effects of beneficial mutations. However, a new read junction was detected between *plaTL055* and *plaTL057* in all sequenced isolates from P5 and P10, suggesting recombination between these plasmids. This pattern implied that encoding all TPA catabolism proteins on a single plasmid reduced burden on the cell, allowing it to grow more quickly.

We also observed an intriguing adaptive pattern emerging in plasmid copy numbers throughout the evolutionary process. Based on *breseq* sequencing analysis, the "fit mean" metric—representing the average read coverage depth across the reference genome—was used as a proxy to estimate plasmid copy numbers by calculating the ratio of plasmid coverage to chromosomal coverage. This analysis revealed variations in the copy numbers of plaTL055 and plaTL057 across different populations (Figure 6), and that plasmid copy numbers appeared to increase over the course of adaptive laboratory evolution. The mechanism underlying this increase was not investigated, but could be related to the recombination events taking place between plaTL055 and plaTL057, or due to a D205V mutation in *pcnB* we observed across all isolates. *pcnB* is believed to play a role in RNAI degradation and the copy number control of CoIE1-related plasmids (He et al., 1993) such as plaTL057. These differences in copy number suggest that the expression levels of the transporter TpaK and the enzyme cascade TphA2A3BA1—encoded by these plasmids—may limit fitness in TPA minimal media.

**Figure 6.**
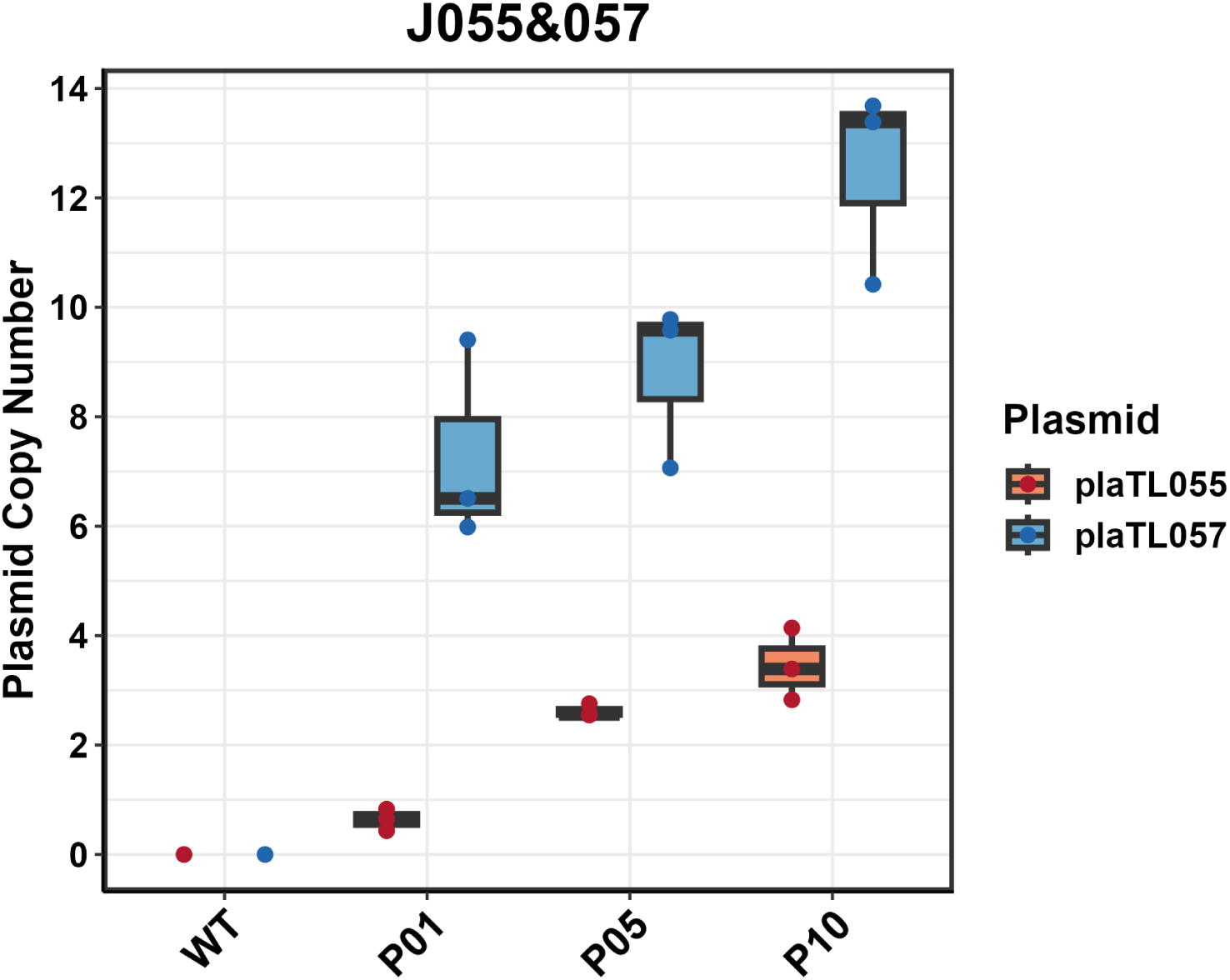
Box plot of J055&057 variants showing plasmid copy number changes across different populations. Sample data are plotted as dots, with boxes indicating quartile regions.

### Tuning TPA Catabolic Genes Using an RBS Library

To address the newly formed junctions between plaTL055 and plaTL057, adjust copy numbers, and alleviate the metabolic burden of maintaining two separate plasmids with different antibiotic requirements, we consolidated these plasmids into a single pET28-based construct while simultaneously constructing RBS libraries for *tpaK* and *tphA2A3BA1*. The goal was to optimize gene expression levels to enhance growth on TPA. RBS libraries were designed using the De Novo DNA RBS library calculator (Farasat et al., 2014; Ng et al., 2015; Reis and Salis, 2020), with details provided in Supplementary File 1.

The J097 RBS library was screened in duplicate in Na₂TP minimal medium. Plasmids from both screenings were extracted and sequenced by Plasmidsaurus, with DADA2 analysis identifying the most frequent RBS for each gene (Supplementary File 3). A summary of these results is provided in Supplementary File 5. We observed that RBS sequences for tphA1I consistently exhibited high TIR values, while the other genes showed more variability in TIR values. However, it is important to note that the TIR values predicted by the De Novo DNA RBS library Calculator may not accurately reflect the real TIR in an operon context. Local sequence context and interactions between RBS sequences in the operon may influence the actual TIR, emphasizing the need for experimental validation of predicted RBS strengths in complex constructs via, for example, proteomics.

From each screening, 10 variants were isolated and further evaluated for growth in Na₂TP minimal medium. All 20 isolates exhibited robust growth, with their calculated intrinsic growth rates significantly higher than those of variants from ALE results, as indicated by a Welch’s t-test p-value of 0.0002. Although the carrying capacities (K) and times to half-maximal carrying capacity (t_mid_) of variants from the RBS library were, on average, higher and shorter than those from the ALE results, the differences were not statistically significant (p_K_ = 0.4248, p_tmid_ = 0.3555). Growth curves for each isolate are shown in Figure 7.

**Figure 7.**
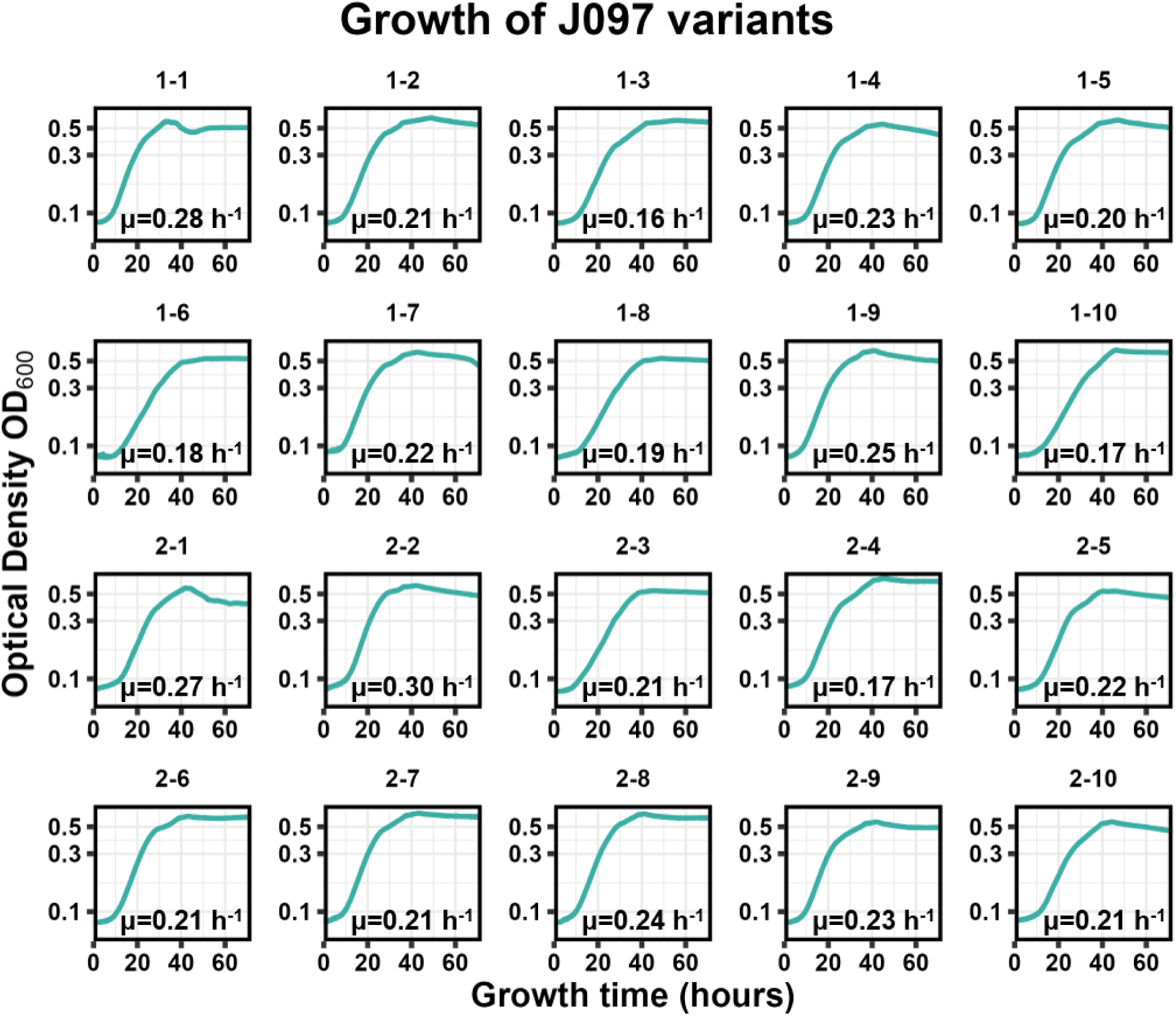
Growth curves of variants isolated from J097 library in Na₂TP minimal media. The y-axis is plotted on a logarithmic scale. Intrinsic growth rates are noted on the figure.

To identify the best-performing variant, scores were assigned to μ, K, and t_mid_, prioritizing variants with the highest μ and K, as well as the shortest t_mid_. Based on the highest overall score, J097-1-9 was selected as the top candidate (plasmid map in Supplementary File 3). J097-1-9 exhibited an intrinsic growth rate of 0.25 h⁻¹, a carrying capacity of 0.47, and took approximately 20 hours to reach half of its carrying capacity. To compare the TIRs for each gene between J097-1-9 and J055&057, we used the De Novo DNA Operon Calculator to predict TIRs. The results showed that most genes in J097-1-9 had lower TIRs, except for tphA2I and tphA1I, which had TIRs 697.42 and 2.65 times higher, respectively, compared to J055&057. The large difference for tphA2I is due to its position as the first expressed gene in the plaTL057 operon, where the originally designed RBS was lower due to operon context. Compared to all variants in the ALE results, J097-1-9 had the highest intrinsic growth rate, demonstrating the benefit of consolidating two plasmids and optimizing translation initiation.

## Discussion and conclusions

The results of this study demonstrate the engineering of *E. coli* for the assimilation of terephthalic acid (TPA), particularly highlighting the successful integration of a TPA transporter and a PCA-producing enzyme cascade. Strain JME3, carrying the PCA degradation pathway, exhibited a notable ability to grow on PCA, which directly informed its suitability for further engineering to assimilate TPA. The selection of JME3 as the chassis strain provided a robust platform for further modifications.

The engineered strain J055&057, which incorporated the TpaK transporter and TphA2A3BA1 enzyme cascade, displayed low growth in TPA minimal medium, while the control strains did not. This finding indicates the functionality of the TpaK transporter and confirms its potential to drive TPA uptake in *E. coli*. Performing 10 rounds of adaptive laboratory evolution successfully enhanced the fitness of J055&057 for growth in TPA minimal medium, shortening the lag phase and improving its growth rate, generating the final population, with a lag phase of 15 hours and a growth rate of 0.22 ± 0.01 h^-1^. While many evolved isolates exhibit the same mutation in genes involved in plasmid copy number, RNA synthesis, and translation, further work is necessary to dissect these mutations and reveal the mechanisms by which they improve growth on TPA.

The observed recombination between plasmids may reflect the fitness trade-offs associated with maintaining the transporter and enzyme genes on separate plasmids. Continued rise in copy number during adaptive laboratory evolution indicated that tuning gene expression was necessary for optimal growth and metabolic performance. The screening of ribosome binding site (RBS) libraries provided valuable insights into optimal expression levels for each gene, culminating in the selection of the best-performing variant, J097-1-9, with an intrinsic growth rate of 0.25 h⁻¹, a carrying capacity of 0.47, and took approximately 20 hours to reach half of its carrying capacity.

However, a comparison of growth in TPA/NaTP minimal medium and PCA minimal medium still revealed that the growth rate J097-1-9 in TPA/Na2TP (0.25 h⁻¹) is significantly slower than JME3 in PCA (0.60 h⁻¹). This observation indicates that some step in the PCA production pathway still limits growth on TPA/Na2TP. It remains unclear whether the transporter efficiency or the conversion of TPA into PCA is the bottleneck. Further modifications to either the transporter or the TPA-to-PCA conversion could potentially improve the growth rate and overall strain performance.

Taken together, this work provides a foundation for further growth improvements on TPA, with potential applications in lignocellulosic biomass conversion, bioremediation of aromatic pollutants, and plastic waste degradation. The integration of transporter and enzyme cascade systems, combined with adaptive evolution, highlights the potential of engineered microbial systems to address critical environmental challenges. By coupling the activities of this cell to PET depolymerization and EG assimilation, an *E. coli-*based biocatalyst could be developed that simultaneously breaks down PET and utilizes its degradation products for biomass regeneration, thereby advancing closed-loop plastic recycling.

## AUTHOR CONTRIBUTIONS

Tianyu Li: Conceptualization (equal); data curation (lead); formal analysis (lead); investigation (lead); methodology (lead); writing – original draft (lead); writing – review and editing (equal). Nathan Crook: Conceptualization (equal); funding acquisition (lead); project administration (lead); supervision (lead); writing – original draft (supporting); writing – review and editing (equal).

## Acknowledgements

We gratefully acknowledge the National Science Foundation (EFMA-2029327) for funding. Dr. Adam Guss for sharing *E. coli* AG978, JME3, and *Rhodococcus jostii* RHA1. We would like to thank Dr. Carly Catella for providing the custom code used in the DADA2 analysis.

## Abbreviations

TPA: terephthalate acid
PET: polyethylene terephthalate
ALE: adaptive laboratory evolution
PCA: protocatechuic acid

## Supplementary

**Figure S1.**
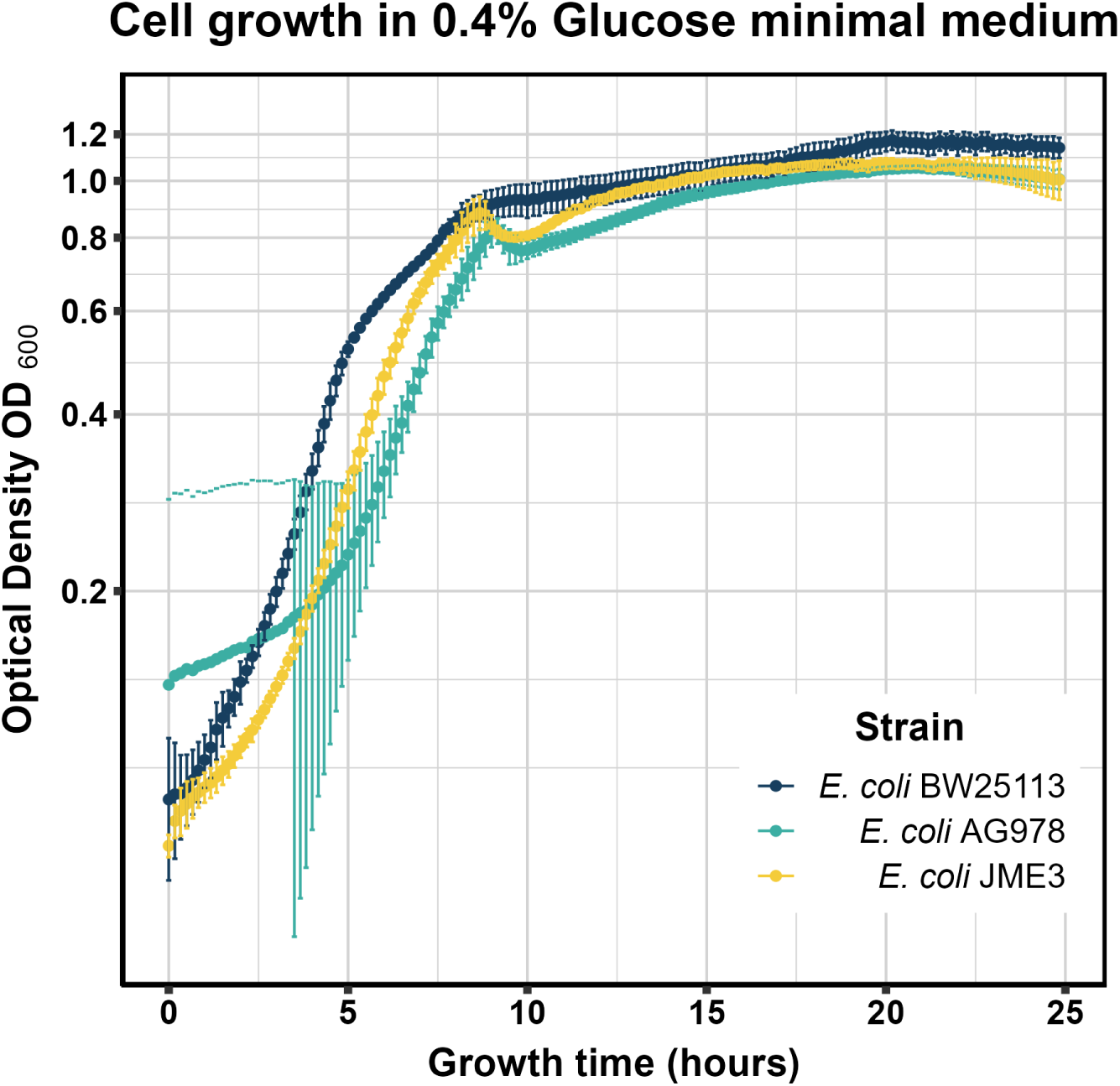
Growth in minimal medium containing 0.4% glucose as the sole carbon source for different *E. coli* strains. Results are presented as mean values, with error bars representing the standard deviation from six technical replicates for each strain.

**Figure S2.**
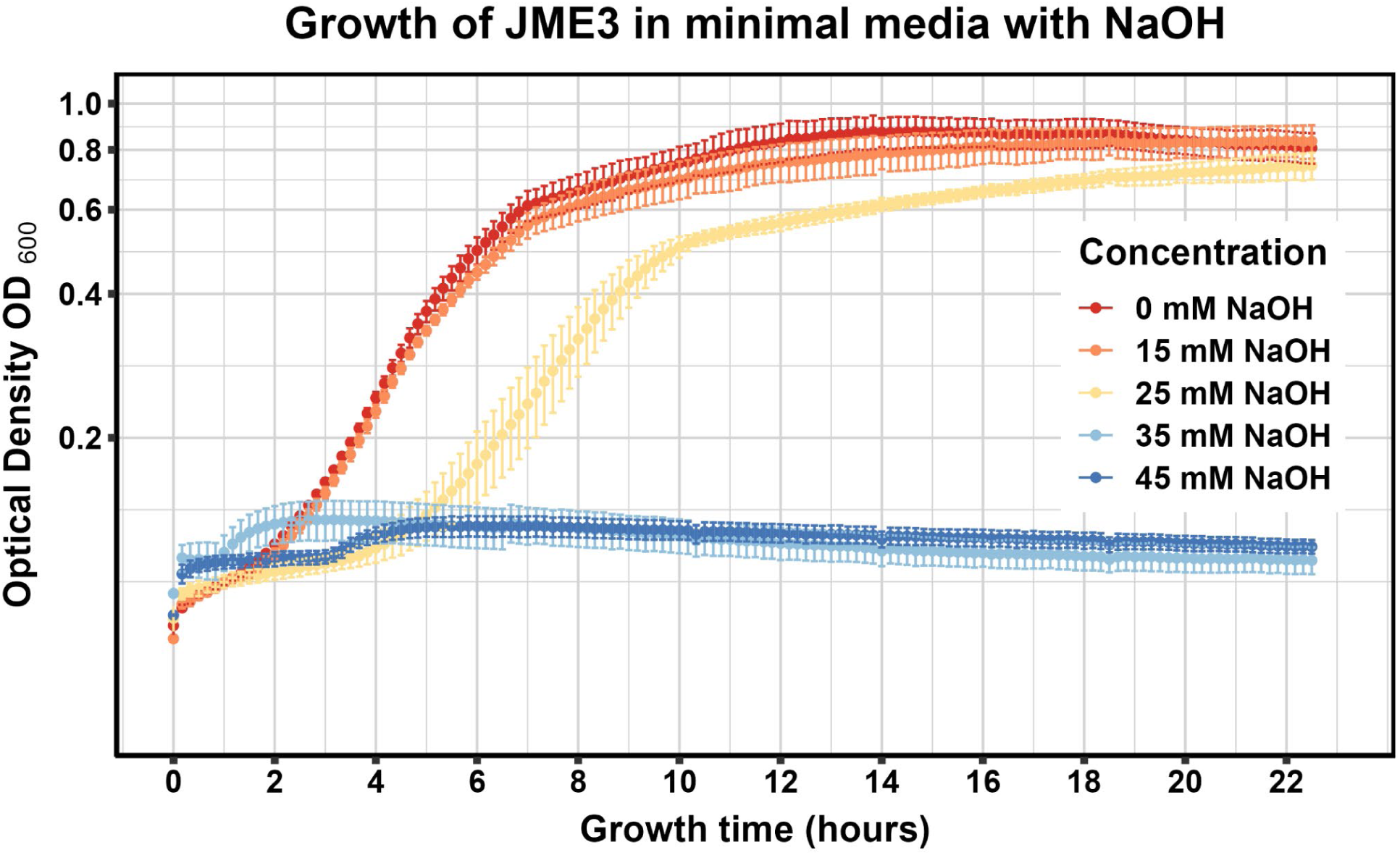
Comparison of growth in minimal medium containing 10 mM glucose for *E. coli* JME3 with varying concentrations of NaOH. Results are presented as mean values, with error bars representing the standard deviation from six technical replicates for each strain.

**Figure S3.**
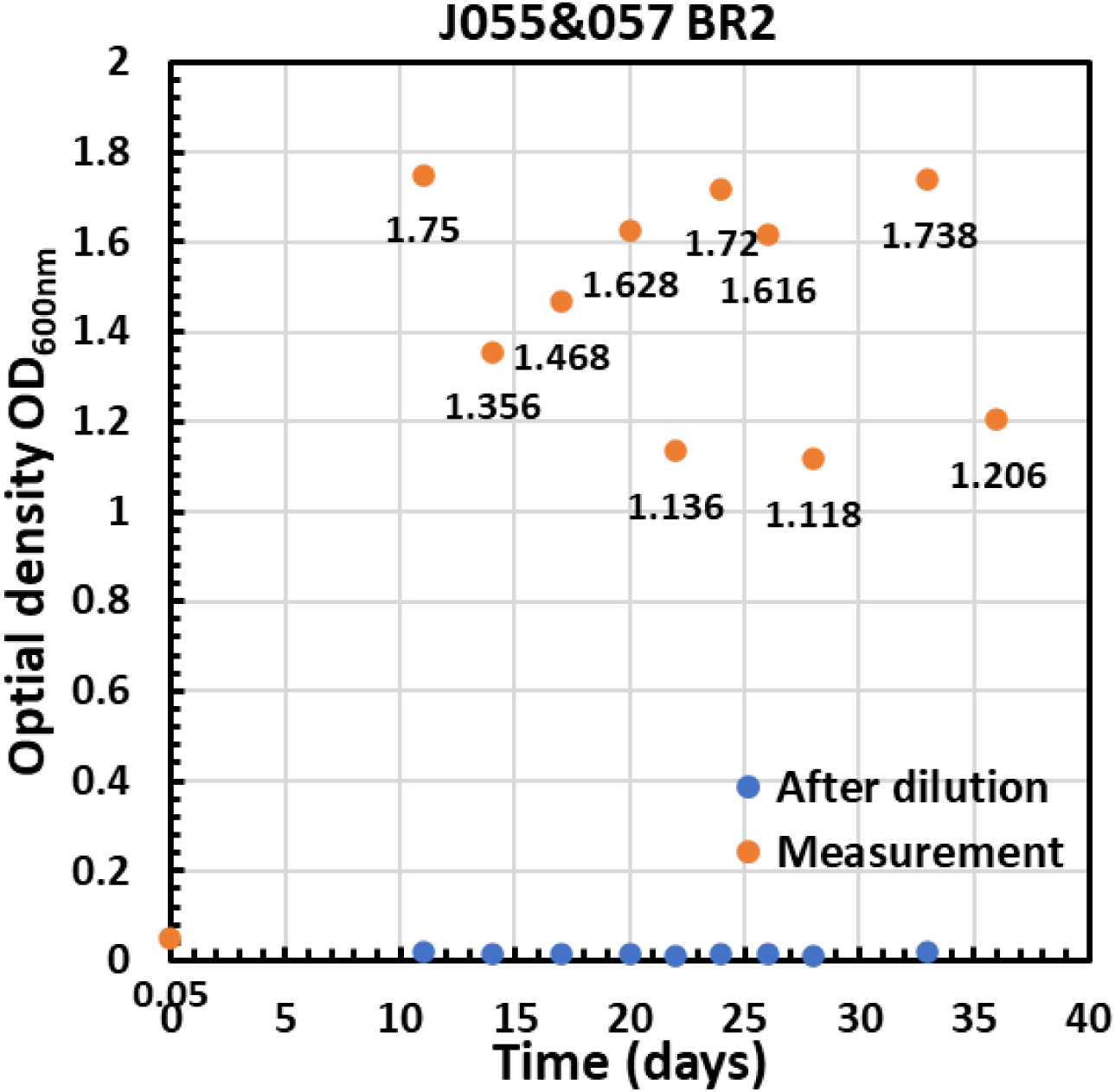
Adaptive evolution of J055&057. Cultures were serially passaged nine times with a 1:100 dilution at each passage. Passing times and OD600 measurements are presented.

**Figure S4.**
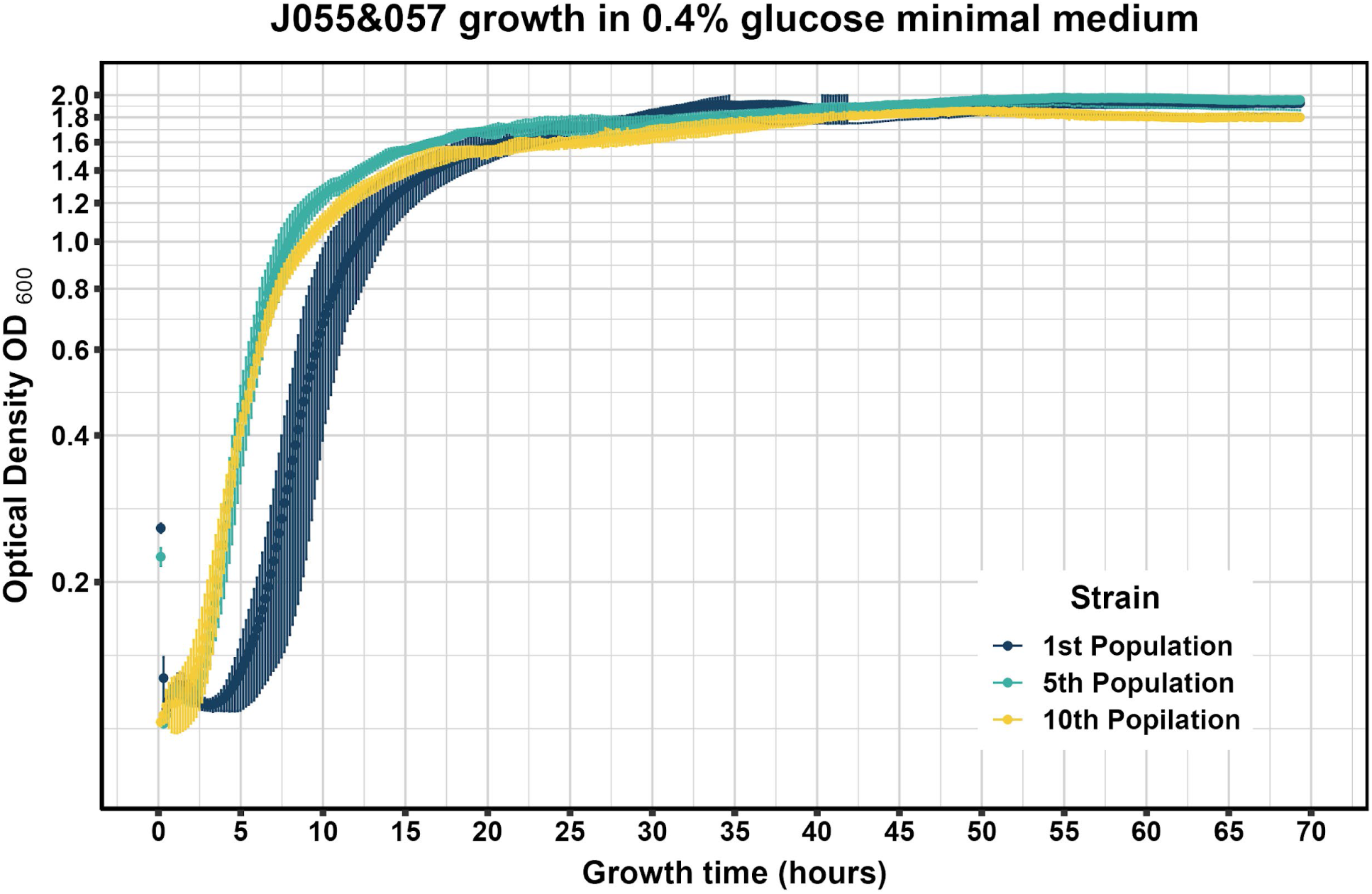
Growth curves of 1st, 5th, and 10th populations of **(A)** J055&057 and **(B)** V055&057 in 0.4% glucose minimal medium. Results are presented as mean values, with error bars representing the standard deviation from six technical replicates for each strain.

**Figure S5.**
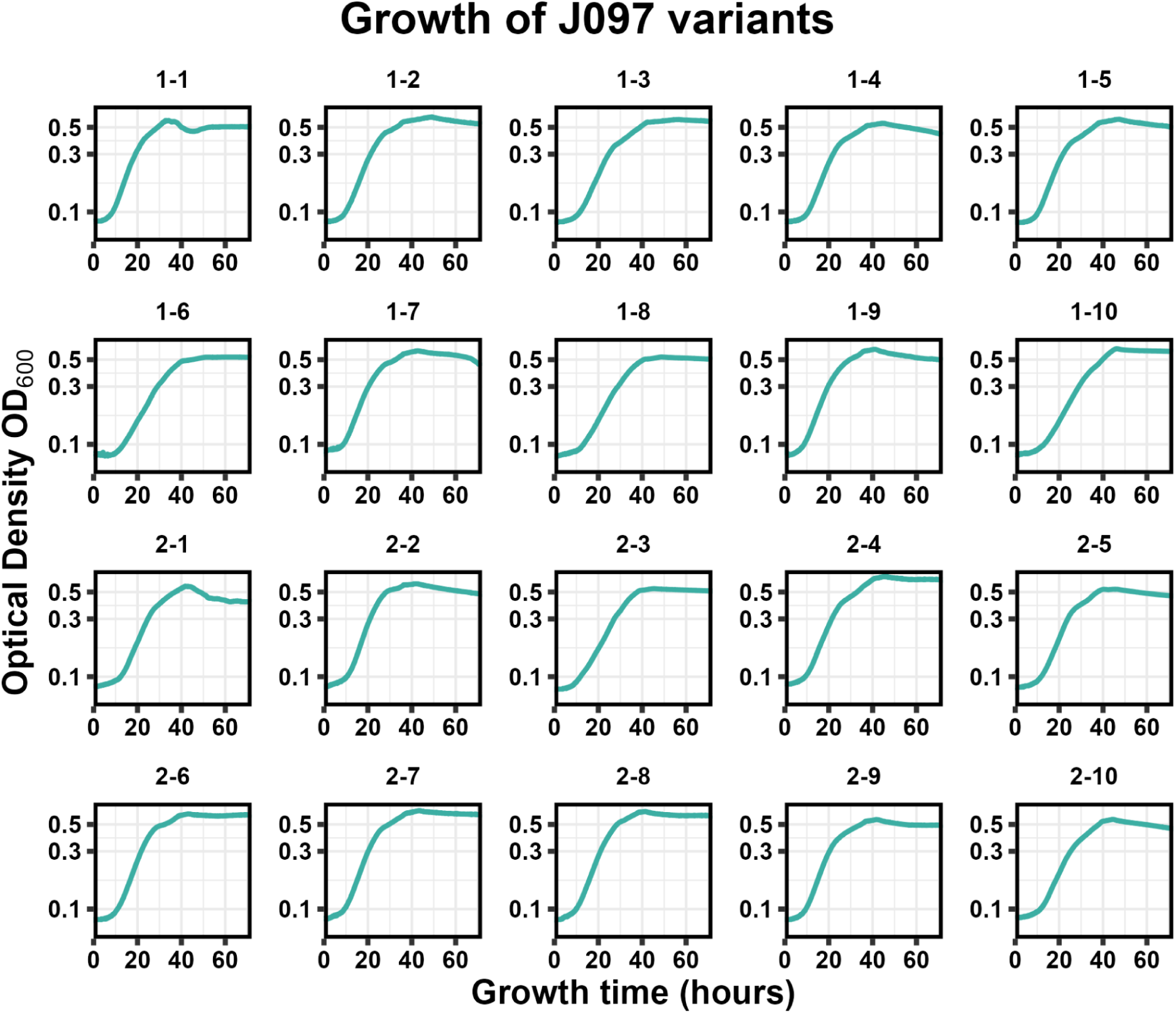
Growth curves of variants isolated from J097 library in Na₂TP minimal media. The y-axis is plotted on a logarithmic scale

## Notes

### Competing Interest Statement

The authors have declared no competing interest.

