## Supplementary File 3 for "Construction of a Heterologous Pathway in *Escherichia coli* for Terephthalate Assimilation": RBSExtraction_2.R

```
# Analyze Plasmidsaurus amplicon sequences
# Identify peptide variants from (NDT)8 library
# Adapted from https://benjjneb.github.io/LRASManuscript/LRASms_Zymo.html
# CMC 6/24/2022

# SETUP
# Need the following
# openssl from https://macappstore.org/openssl/
# Bioconductor packages from https://www.bioconductor.org/install/
# Load libraries, setup paths, prepare environment:
library(dada2);packageVersion("dada2")
library(Biostrings);packageVersion("Biostrings")
library(ShortRead);packageVersion("ShortRead")
library(ggplot2);packageVersion("ggplot2")
library(reshape2);packageVersion("reshape2")
library(RColorBrewer);packageVersion("RColorBrewer")
library(ORFik);packageVersion("ORFik")
library(Rsamtools);packageVersion("Rsamtools")
library(seqTools);packageVersion("seqTools")

path <- "C:/Users/69467/Desktop/Li_P6B_raw_reads" # CHANGE ME to location of the fastq file
path.out <- "Figures/" 
path.rds <- "RDS/" 
rfn <-"Li_P6B_11_plaTL118-1-pooled.fastq" #root file name

fn <- file.path(path, rfn)
# F_epPCR <- "cgtccggcgtagaggatcgaga" # primers used for amplification not present in sequences
# R_epPCR <- "gcagggaagcgctagagatcct" # primers used for amplification not present in sequences
# F_flank <- "cacacataaacaaac" # mock primer adjacent to region of interest
# R_flank <- "gaaaaacggta" # mock primer adjacent to region of interest

#TPA pathway primers
F_flank <- "cgtccggcgtagaggatcgaga" # mock primer adjacent to region of interest (crops around peptide)
R_flank <- "gcagggaagcgctagagatcct" # mock primer adjacent to region of interest (crops around peptide)
rc <- dada2:::rc
theme_set(theme_bw())

# REMOVE PRIMERS AND FILTER
# Remove primers and orient reads:
npfn <-paste("no_primer",rfn, sep = "_") #no primer files names
nop <- file.path(path, "noprimers", basename(npfn))
prim <- removePrimers(fn, nop, primer.fwd=F_flank, primer.rev=dada2:::rc(R_flank), allow.indels=TRUE, trim.fwd=TRUE, trim.rev=TRUE, orient=TRUE, verbose=TRUE,compress=FALSE)
hist(nchar(getSequences(nop)), 100)
# filter
ffn <-paste("filtered",rfn, sep = "_") #filtered files names
filt<-file.path(path, "noprimers","filtered", basename(ffn))
track <- fastqFilter(nop, filt, minQ=0,  minLen=3000,maxLen=4000, trimLeft=0, maxN=0, rm.phix=FALSE, verbose=TRUE, compress=FALSE) 
# trimLeft implemented to remove remaining primer
# minLen=6 to prevent code from breaking at error step (min length > k-mer = 5)
hist(nchar(getSequences(filt)), 100)

# RUN DADA2
# Dereplicate
drp <- derepFastq(filt, verbose=TRUE)
# Learn errors
#err <- learnErrors(drp, BAND_SIZE=32, multithread=TRUE, errorEstimationFunction=dada2:::PacBioErrfun) # 10s of seconds
err <- learnErrors(drp, BAND_SIZE=32, multithread=TRUE) # 10s of seconds, removed PacBioErrfun
plotErrors(err)
# Denoise
dd <- dada(drp, err=err, BAND_SIZE=32, multithread=TRUE) # seconds
dd
# Read tracking
cbind(ccs=prim[,1], primers=prim[,2], filtered=track[[2]], denoised=sum(dd$denoised))
# Inspect clustering data.frame
dd$clustering[,-1]

# Get Uniques
ufn <-paste("uni",rfn, sep = "_") #uniqe file name 
uniques <-- getUniques(dd)
fna <- file.path(path, ufn)
Uniques <- file.path(path, "noprimers", "filtered", "uniques", basename(fna))
uniquesToFasta(uniques, Uniques)


# VIEW RAW
# dada2 compression only works well when there are many amplicons for a given sequence
# filteres out "noisey" low abundance reads
qfn <-paste("quantify",rfn, sep = "_") #quantify files names
uni <- getUniques(drp) # uniques prior to dada compression
fna1 <- file.path(path, qfn)
Uni <- file.path(path, "noprimers", "filtered", "uniques", basename(fna1))
uniquesToFasta(uni, Uni)


######tpaK RBS primers
F_flank <- "ataacaatttcacac" # mock primer adjacent to region of interest (crops around peptide)
R_flank <- "atggtgctaaggacat" # mock primer adjacent to region of interest (crops around peptide)
rc <- dada2:::rc
theme_set(theme_bw())

# REMOVE PRIMERS AND FILTER
# Remove primers and orient reads:
tpaK_npfn <-paste("tpaK_no_primer",rfn, sep = "_") #no primer files names
nop <- file.path(path, "noprimers", basename(tpaK_npfn))
prim <- removePrimers(fn, nop, primer.fwd=F_flank, primer.rev=dada2:::rc(R_flank), allow.indels=TRUE, trim.fwd=TRUE, trim.rev=TRUE, orient=TRUE, verbose=TRUE,compress=FALSE)
hist(nchar(getSequences(nop)), 100)
# filter
tpaK_ffn <-paste("tpaK_filtered",rfn, sep = "_") #filtered files names
filt<-file.path(path, "noprimers","filtered", basename(tpaK_ffn))
track <- fastqFilter(nop, filt, minQ=3, minLen=15, maxLen=15, trimLeft=0, maxN=0, rm.phix=FALSE, verbose=TRUE, compress=FALSE) 
# trimLeft implemented to remove remaining primer
# minLen=6 to prevent code from breaking at error step (min length > k-mer = 5)
hist(nchar(getSequences(filt)), 100)

# RUN DADA2
# Dereplicate
drp <- derepFastq(filt, verbose=TRUE)
# Learn errors
#err <- learnErrors(drp, BAND_SIZE=32, multithread=TRUE, errorEstimationFunction=dada2:::PacBioErrfun) # 10s of seconds
err <- learnErrors(drp, BAND_SIZE=32, multithread=TRUE) # 10s of seconds, removed PacBioErrfun
plotErrors(err)
# Denoise
dd <- dada(drp, err=err, BAND_SIZE=32, multithread=TRUE) # seconds
dd
# Read tracking
cbind(ccs=prim[,1], primers=prim[,2], filtered=track[[2]], denoised=sum(dd$denoised))
# Inspect clustering data.frame
dd$clustering[,-1]

# Get Uniques
tpaK_ufn <-paste("tpaK_uni",rfn, sep = "_") #uniqe file name 
uniques <-- getUniques(dd)
fna <- file.path(path, tpaK_ufn)
Uniques <- file.path(path, "noprimers", "filtered", "uniques", basename(fna))
uniquesToFasta(uniques, Uniques)


# VIEW RAW
# dada2 compression only works well when there are many amplicons for a given sequence
# filteres out "noisey" low abundance reads
tpaK_qfn <-paste("tpaK_quantify",rfn, sep = "_") #quantify files names
uni <- getUniques(drp) # uniques prior to dada compression
fna1 <- file.path(path, tpaK_qfn)
Uni <- file.path(path, "noprimers", "filtered", "uniques", basename(fna1))
uniquesToFasta(uni, Uni)

######tphA2I RBS primers
F_flank <- "attcgctcgcccgcaagtag" # mock primer adjacent to region of interest (crops around peptide)
R_flank <- "tggatgatggattcttgcat" # mock primer adjacent to region of interest (crops around peptide)
rc <- dada2:::rc
theme_set(theme_bw())

# REMOVE PRIMERS AND FILTER
# Remove primers and orient reads:
tphA2I_npfn <-paste("tphA2I_no_primer",rfn, sep = "_") #no primer files names
nop <- file.path(path, "noprimers", basename(tphA2I_npfn))
prim <- removePrimers(fn, nop, primer.fwd=F_flank, primer.rev=dada2:::rc(R_flank), allow.indels=TRUE, trim.fwd=TRUE, trim.rev=TRUE, orient=TRUE, verbose=TRUE,compress=FALSE)
hist(nchar(getSequences(nop)), 100)
# filter
tphA2I_ffn <-paste("tphA2I_filtered",rfn, sep = "_") #filtered files names
filt<-file.path(path, "noprimers","filtered", basename(tphA2I_ffn))
track <- fastqFilter(nop, filt, minQ=3, minLen=17, maxLen=17, trimLeft=0, maxN=0, rm.phix=FALSE, verbose=TRUE, compress=FALSE) 
# trimLeft implemented to remove remaining primer
# minLen=6 to prevent code from breaking at error step (min length > k-mer = 5)
hist(nchar(getSequences(filt)), 100)

# RUN DADA2
# Dereplicate
drp <- derepFastq(filt, verbose=TRUE)
# Learn errors
#err <- learnErrors(drp, BAND_SIZE=32, multithread=TRUE, errorEstimationFunction=dada2:::PacBioErrfun) # 10s of seconds
err <- learnErrors(drp, BAND_SIZE=32, multithread=TRUE) # 10s of seconds, removed PacBioErrfun
plotErrors(err)
# Denoise
dd <- dada(drp, err=err, BAND_SIZE=32, multithread=TRUE) # seconds
dd
# Read tracking
cbind(ccs=prim[,1], primers=prim[,2], filtered=track[[2]], denoised=sum(dd$denoised))
# Inspect clustering data.frame
dd$clustering[,-1]

# Get Uniques
tphA2I_ufn <-paste("tphA2I_uni",rfn, sep = "_") #uniqe file name 
uniques <-- getUniques(dd)
fna <- file.path(path, tphA2I_ufn)
Uniques <- file.path(path, "noprimers", "filtered", "uniques", basename(fna))
uniquesToFasta(uniques, Uniques)


# VIEW RAW
# dada2 compression only works well when there are many amplicons for a given sequence
# filteres out "noisey" low abundance reads
tphA2I_qfn <-paste("tphA2I_quantify",rfn, sep = "_") #quantify files names
uni <- getUniques(drp) # uniques prior to dada compression
fna1 <- file.path(path, tphA2I_qfn)
Uni <- file.path(path, "noprimers", "filtered", "uniques", basename(fna1))
uniquesToFasta(uni, Uni)


######tphA3II RBS primers
F_flank <- "gcatatgggacaggagatgcaagcatga" # mock primer adjacent to region of interest (crops around peptide)
R_flank <- "gcgatttgaatttcattgatcat" # mock primer adjacent to region of interest (crops around peptide)
rc <- dada2:::rc
theme_set(theme_bw())

# REMOVE PRIMERS AND FILTER
# Remove primers and orient reads:
tphA3II_npfn <-paste("tphA3II_no_primer",rfn, sep = "_") #no primer files names
nop <- file.path(path, "noprimers", basename(tphA3II_npfn))
prim <- removePrimers(fn, nop, primer.fwd=F_flank, primer.rev=dada2:::rc(R_flank), allow.indels=TRUE, trim.fwd=TRUE, trim.rev=TRUE, orient=TRUE, verbose=TRUE,compress=FALSE)
hist(nchar(getSequences(nop)), 100)
# filter
tphA3II_ffn <-paste("tphA3II_filtered",rfn, sep = "_") #filtered files names
filt<-file.path(path, "noprimers","filtered", basename(tphA3II_ffn))
track <- fastqFilter(nop, filt, minQ=3, minLen=15, maxLen=15, trimLeft=0, maxN=0, rm.phix=FALSE, verbose=TRUE, compress=FALSE) 
# trimLeft implemented to remove remaining primer
# minLen=6 to prevent code from breaking at error step (min length > k-mer = 5)
hist(nchar(getSequences(filt)), 100)

# RUN DADA2
# Dereplicate
drp <- derepFastq(filt, verbose=TRUE)
# Learn errors
#err <- learnErrors(drp, BAND_SIZE=32, multithread=TRUE, errorEstimationFunction=dada2:::PacBioErrfun) # 10s of seconds
err <- learnErrors(drp, BAND_SIZE=32, multithread=TRUE) # 10s of seconds, removed PacBioErrfun
plotErrors(err)
# Denoise
dd <- dada(drp, err=err, BAND_SIZE=32, multithread=TRUE) # seconds
dd
# Read tracking
cbind(ccs=prim[,1], primers=prim[,2], filtered=track[[2]], denoised=sum(dd$denoised))
# Inspect clustering data.frame
dd$clustering[,-1]

# Get Uniques
tphA3II_ufn <-paste("tphA3II_uni",rfn, sep = "_") #uniqe file name 
uniques <-- getUniques(dd)
fna <- file.path(path, tphA3II_ufn)
Uniques <- file.path(path, "noprimers", "filtered", "uniques", basename(fna))
uniquesToFasta(uniques, Uniques)


# VIEW RAW
# dada2 compression only works well when there are many amplicons for a given sequence
# filteres out "noisey" low abundance reads
tphA3II_qfn <-paste("tphA3II_quantify",rfn, sep = "_") #quantify files names
uni <- getUniques(drp) # uniques prior to dada compression
fna1 <- file.path(path, tphA3II_qfn)
Uni <- file.path(path, "noprimers", "filtered", "uniques", basename(fna1))
uniquesToFasta(uni, Uni)

######tphBI RBS primers
F_flank <- "tgatggcattgccgctatga" # mock primer adjacent to region of interest (crops around peptide)
R_flank <- "atggccaaagccaatctacggtgcactattgtcat" # mock primer adjacent to region of interest (crops around peptide)
rc <- dada2:::rc
theme_set(theme_bw())

# REMOVE PRIMERS AND FILTER
# Remove primers and orient reads:
tphBI_npfn <-paste("tphBI_no_primer",rfn, sep = "_") #no primer files names
nop <- file.path(path, "noprimers", basename(tphBI_npfn))
prim <- removePrimers(fn, nop, primer.fwd=F_flank, primer.rev=dada2:::rc(R_flank), allow.indels=TRUE, trim.fwd=TRUE, trim.rev=TRUE, orient=TRUE, verbose=TRUE,compress=FALSE)
hist(nchar(getSequences(nop)), 100)
# filter
tphBI_ffn <-paste("tphBI_filtered",rfn, sep = "_") #filtered files names
filt<-file.path(path, "noprimers","filtered", basename(tphBI_ffn))
track <- fastqFilter(nop, filt, minQ=3, minLen=15, maxLen=15, trimLeft=0, maxN=0, rm.phix=FALSE, verbose=TRUE, compress=FALSE) 
# trimLeft implemented to remove remaining primer
# minLen=6 to prevent code from breaking at error step (min length > k-mer = 5)
hist(nchar(getSequences(filt)), 100)

# RUN DADA2
# Dereplicate
drp <- derepFastq(filt, verbose=TRUE)

# Denoise
dd <- dada(drp, err=err, BAND_SIZE=32, multithread=TRUE) # seconds
dd
# Read tracking
cbind(ccs=prim[,1], primers=prim[,2], filtered=track[[2]], denoised=sum(dd$denoised))
# Inspect clustering data.frame
dd$clustering[,-1]

# Get Uniques
tphBI_ufn <-paste("tphBI_uni",rfn, sep = "_") #uniqe file name 
uniques <-- getUniques(dd)
fna <- file.path(path, tphBI_ufn)
Uniques <- file.path(path, "noprimers", "filtered", "uniques", basename(fna))
uniquesToFasta(uniques, Uniques)


# VIEW RAW
# dada2 compression only works well when there are many amplicons for a given sequence
# filteres out "noisey" low abundance reads
tphBI_qfn <-paste("tphBI_quantify",rfn, sep = "_") #quantify files names
uni <- getUniques(drp) # uniques prior to dada compression
fna1 <- file.path(path, tphBI_qfn)
Uni <- file.path(path, "noprimers", "filtered", "uniques", basename(fna1))
uniquesToFasta(uni, Uni)

######tphA1I RBS primers
F_flank <- "cgccctgctcggcgcccaaccgggctga" # mock primer adjacent to region of interest (crops around peptide)
R_flank <- "gtcgtggatatggatctggtggttcat" # mock primer adjacent to region of interest (crops around peptide)
rc <- dada2:::rc
theme_set(theme_bw())

# REMOVE PRIMERS AND FILTER
# Remove primers and orient reads:
tphA1I_npfn <-paste("tphA1I_no_primer",rfn, sep = "_") #no primer files names
nop <- file.path(path, "noprimers", basename(tphA1I_npfn))
prim <- removePrimers(fn, nop, primer.fwd=F_flank, primer.rev=dada2:::rc(R_flank), allow.indels=TRUE, trim.fwd=TRUE, trim.rev=TRUE, orient=TRUE, verbose=TRUE,compress=FALSE)
hist(nchar(getSequences(nop)), 100)
# filter
tphA1I_ffn <-paste("tphA1I_filtered",rfn, sep = "_") #filtered files names
filt<-file.path(path, "noprimers","filtered", basename(tphA1I_ffn))
track <- fastqFilter(nop, filt, minQ=3, minLen=15, maxLen=15, trimLeft=0, maxN=0, rm.phix=FALSE, verbose=TRUE, compress=FALSE) 
# trimLeft implemented to remove remaining primer
# minLen=6 to prevent code from breaking at error step (min length > k-mer = 5)
hist(nchar(getSequences(filt)), 100)

# RUN DADA2
# Dereplicate
drp <- derepFastq(filt, verbose=TRUE)
# Learn errors
#err <- learnErrors(drp, BAND_SIZE=32, multithread=TRUE, errorEstimationFunction=dada2:::PacBioErrfun) # 10s of seconds
err <- learnErrors(drp, BAND_SIZE=32, multithread=TRUE) # 10s of seconds, removed PacBioErrfun
plotErrors(err)
# Denoise
dd <- dada(drp, err=err, BAND_SIZE=32, multithread=TRUE) # seconds
dd
# Read tracking
cbind(ccs=prim[,1], primers=prim[,2], filtered=track[[2]], denoised=sum(dd$denoised))
# Inspect clustering data.frame
dd$clustering[,-1]

# Get Uniques
tphA1I_ufn <-paste("tphA1I_uni",rfn, sep = "_") #uniqe file name 
uniques <-- getUniques(dd)
fna <- file.path(path, tphA1I_ufn)
Uniques <- file.path(path, "noprimers", "filtered", "uniques", basename(fna))
uniquesToFasta(uniques, Uniques)


# VIEW RAW
# dada2 compression only works well when there are many amplicons for a given sequence
# filteres out "noisey" low abundance reads
tphA1I_qfn <-paste("tphA1I_quantify",rfn, sep = "_") #quantify files names
uni <- getUniques(drp) # uniques prior to dada compression
fna1 <- file.path(path, tphA1I_qfn)
Uni <- file.path(path, "noprimers", "filtered", "uniques", basename(fna1))
uniquesToFasta(uni, Uni)
```
